# Model-free machine learning-based 3D single molecule localisation microscopy

**DOI:** 10.1101/2024.10.14.618179

**Authors:** Miguel A. Boland, Jonathan P. E. Lightley, Edwin Garcia, Sunil Kumar, Chris Dunsby, Seth Flaxman, Mark A. A. Neil, Paul M. W. French, Edward A. K. Cohen

**Author notes:** **Correspondence** Miguel A. Boland. Department of Mathematics, Imperial College London, South Kensington, London, SW7 2AZ, U.K.

## Abstract

Single molecule localisation microscopy (SMLM) can provide two-dimensional super-resolved image data from conventional fluorescence microscopes, while three dimensional (3D) SMLM usually involves a modification of the microscope, e.g. to engineer a predictable axial variation in the point spread function. Here we demonstrate a 3D SMLM approach (we call *“easyZloc”*) utilising a lightweight Convolutional Neural Network that is generally applicable, including with “standard” (unmodified) fluorescence microscopes, and which we consider may be practically useful in a high throughput SMLM workflow. We demonstrate the reconstruction of nuclear pore complexes with comparable performance to previously reported methods but with a significant reduction in computational power and execution time. 3D reconstructions of the nuclear envelope and a tubulin sample over a larger axial range are also shown.

## 1 INTRODUCTION

Single Molecule Localisation Microscopy (SMLM) is used to describe a range of imaging techniques to achieve spatial resolution beyond the diffraction-limit in fluorescence microscopes, as reviewed in, e.g.,^1–3^. These techniques enable super-resolved imaging of fluorescently labelled nanoscale structures with transverse spatial resolution approaching that of electron microscopy. This is achieved by acquiring a sequence of images in which only a sparse subset of fluorophores emit light during each camera acquisition, such that their point spread functions can be distinguished. Thus, the location of each fluorophore in the image plane can be determined, e.g., by centroiding the discrete images of individual emitters, to achieve an enhanced precision that depends on the number of photons detected. The most commonly used SMLM techniques include Photo Activated Localisation Microscopy (PALM)^4^, Stochastic Optical Reconstruction Microscopy (STORM)^5^ – particularly implementations based on direct photoswitching of conventional fluorophores such as dSTORM^6^ and GSD-IM^7^ - and techniques based on transient binding of diffusing fluorophores such as Points Accumulation for Imaging in Nanoscale Topography (PAINT)^8^ and DNA-PAINT^9^. In general, SMLM techniques are relatively straightforward to implement on existing epifluorescence microscopes^10–12^; we and others have demonstrated low-cost SMLM instruments using multimode diode laser excitation with inexpensive CMOS cameras that can help widen access to super-resolved microscopy.

It is also possible to localise sparse emitters in the axial (Z) direction to realise 3D SMLM by analysing the shape of the image of each emitter where the PSF varies with Z. This is most frequently realised in microscopes that have been modified to exhibit significant astigmatism, e.g., through the insertion of a cylindrical lens in the imaging path^13^, as previously used to axially localise quantum dots^14^. Alternatively, the microscope PSF can be explicitly engineered, e.g. using wavefront-shaping techniques, to provide an extended axial range of Z localisation. The first implementation of this wavefront-shaping approach was the double helix PSF^15^, and this has subsequently been extended to further engineered PSF, e.g.^16^. A third approach involves imaging the sparse emitters at two or more different focal planes, e.g.^17^, and comparing the multiple PSFs corresponding to each detected emitter. These and other 3D SMLM techniques entail modification of the fluorescence microscope with varying degrees of complexity and cost; the simplest approach using cylindrical lens-induced astigmatism provides high axial precision but is limited to a relatively small (∼500 nm) range.

To further widen access to SMLM and other microscopy techniques, we are developing a cost-effective open-source microscope platform that utilises a modular microscope stand (*“openFrame”*)^18^ to implement our low cost *“easySTORM”* approach^10^ to SMLM, and we are working towards a practical open-source cost-effective platform for automated SMLM of samples arrayed in multiwell plates^10,18^. As part of our efforts to minimise cost, while maximising access and flexibility in these instruments, we explore utilising machine learning to determine the Z-localisation of emitting fluorophores with no modification of the fluorescence microscope, i.e., directly from the image data acquired for 2D SMLM exploiting only the microscope’s intrinsic aberration. Our approach is to obtain “ground truth” training data by acquiring calibration Z-stacks of images of sub-resolution fluorescent beads in order to map the axial and lateral variation of the microscope’s intrinsic PSF across its field of view. This approach, which we term *easyZloc*, makes no assumptions about the instrument and can, in principle, be applied to any microscope configuration, i.e., any combination of microscope optics and camera, as long as a “calibration” Z-stack of sub-resolution fluorescent beads is acquired before an imaging session.

We note that Z-localisation using the microscope’s intrinsic axial variation of experimental PSF has previously been explored using an analytic model-based fitting approach with 2D SMLM data^19,20^, and that several groups have utilised deep learning for 3D SMLM using astigmatic or engineered PSF, e.g.^21–24^. Most of this prior work uses convolutional neural networks (CNN) to determine the localisation of each emitter in three dimensions and typically requires significant computational resources, often limiting them to relatively small fields of view or requiring significant computation times that would not be practical for some applications, particularly automated multiwell plate SMLM where we aim to acquire and process SMLM data from hundreds of large (>100 µm) fields of view (FOV).

Prior work based on phase-retrieval methods leverage Zernike polynomials to model the microscope PSF, for example fitting a model to a calibration image of fluorescent beads^25^ and generating simulated noise-free samples to fit a Maximum Likelihood Estimate (MLE) of the localisation’s depth. In-situ PSF Retrieval (INSPR)^23^ extended this approach by constructing the Zernike model directly from experimental data whilst iteratively localising the dataset. These methods assume a uniform PSF model across the FOV of the microscope and were demonstrated with microscopes engineered to present PSFs with significant axial variation. An alternative paradigm for 3D localisation methodologies relies on empirical models of the PSF that are not constrained by an underlying Zernike polynomial. For example, the use of a 3D cubic spline and MLE to model a PSF was demonstrated on SMLM data from microscopes with astigmatic and saddle point PSFs^26^. This approach yielded an improved localisation accuracy compared to earlier Gaussian based models and should be applicable with any experimental PSF as no assumptions are made concerning a theoretical model of the PSF. This approach was further refined into a GPU-accelerated implementation of many different PSF modalities^19^. However, these approaches also assume a constant PSF across the FOV and could potentially be compromised in microscopes with a significant lateral variation of the PSF aberrations across the FOV, e.g., due to spherical or other aberrations.

Further approaches such as *DeepLoc*^27^, *DeepSTORM3D*^28^ and *DECODE*^22^ have also leveraged deep learning for whole image localisation, for example using cubic spline models of an experimental PSF to generate simulated training data^22^. To date, these methods have modelled laterally invariant PSFs^27^, relatively complex (and expensive to implement) tetrapod PSFs^28^ or highly astigmatic / double-helix PSFs^22^. To the best of our knowledge only one deep learning-based method, *FD-DeepLoc*^21^, has accounted for lateral variations in PSF aberrations, implementing a vectorial Zernike model fitted to experimental PSF data, but this was demonstrated on optical systems with a strong astigmatism induced by a cylindrical lens or a tetrapod PSF.

Our approach is to realise 3D SMLM with minimal perturbations to the microscope hardware, leveraging existing SMLM workflows and minimal computational requirements. We approach this by using established (2D) SMLM workflows such as *ThunderSTORM*^29^ or *PICASSO*^30^ for transverse localisation of the emitters and then determining their axial localisation using a rapid CNN-based approach, as illustrated in Figure 1. To the best of our knowledge, this is the first attempt to determine Z-localisation on a “standard” fluorescence microscope (i.e., with no engineered astigmatism) using deep learning trained with experimental SMLM calibration data acquired on the same instrument. It should be applicable with a wide range of experimental PSF and does not require a theoretical basis for the PSF, e.g. from fitting to cubic splines or a Zernike-based PSF model.

**FIGURE 1:**
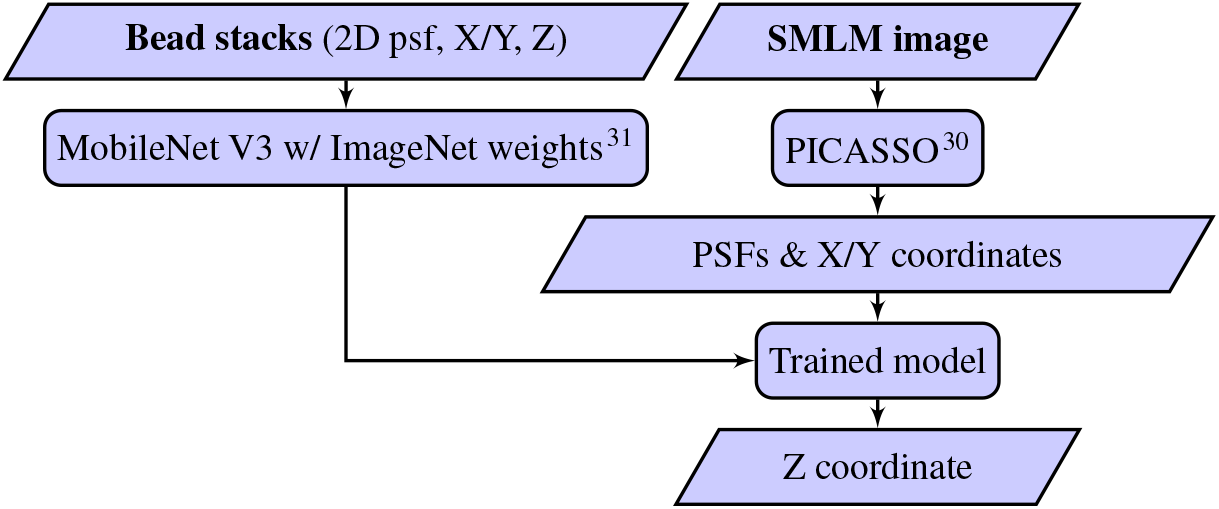
Application of *easyZloc* on experimental data

We leverage the robustness of MobileNet v3^32^ to map the experimental 3D spatially varying PSF, including any optical aberrations, directly from a calibration image of fluorescent sub-resolution (TetraSpeck) beads (Section 2.2, Section 2.4). MobileNet v3 was selected as it is the latest version of a computationally efficient high performance convolutional neural network. Lateral single molecule localisation is first undertaken using a separate software package (*PICASSO*)^30^ to transform the acquired SMLM data into a 2D localisation table and a set of extracted 2D images of localisations. These are then localised in Z using our trained model. We demonstrate this approach can provide a relatively fast and robust solution to achieve 3D localisation in almost any SMLM microscope, without modification to engineer the PSF and with modest computational requirements. When imaging fluorescently labelled nuclear pore proteins (using *easySTORM* on a standard epifluorescence microscope with HiLo illumination), we were able to resolve structures axially separated by ∼ 50 nm. In a first attempt to benchmark our approach, we also applied *FD-DeepLoc* and *DECODE* to the same SMLM data – albeit noting that these software tools were developed and evaluated using microscopes exhibiting significant astigmatism from engineered PSF – and found that our MobileNet v3-based *easyZloc* approach provided at least comparable reconstructions of a (minimally astigmatic) nuclear pore SMLM data set, while requiring significantly less computational time.

## 2 METHODS

### 2.1 Single molecule localisation microscope

A commercial epifluorescence microscope frame (Axiovert 200, Zeiss) was used to acquire the *easySTORM* SMLM data presented in this paper. As outlined in^10^, this instrument utilises a vibrating multimode optical fibre to deliver excitation light from multimode laser diodes within a commercial multiline laserbank (Multiline laserbank, Cairn Research Ltd). Using Köhler illumination, this provides uniform illumination over a large (>120 µm diameter) FOV for *easySTORM*^10^. Sample excitation could be switched between HiLo^33^ and epi-illumination using an excitation coupling unit positioned in the back port of the commercial frame (OptoTIRF; Cairn Research Ltd). A custom-built hardware based optical autofocus module^18^ was used to maintain focus throughout the SMLM acquisitions. For the data presented in this paper, an sCMOS camera (Photometrics Prime 95B) was used to acquire both the training calibration data and the fluorescent SMLM data. This microscope was fitted with a piezoelectric Z-stage (NanoScan NZ100, Prior) and a motorized XY stage (MS-2000, Applied Scientific Instruments) to control axial and lateral movement. The hardware was all controlled using *µManager* 2.0^34^.

### 2.2 Calibration sample preparation and imaging

Bead samples to acquire the calibration training data were prepared as follows: diluted 1 : 10^7^ per millilitre fluorescent polystyrene beads with a diameter of 20 nm, excitation/emission: 625/645 nm (F8782-Invitrogen FluoSpheres™ Carboxylate-Modified Microspheres) were attached to a glass 8-well chamber slide 1.5H (80827 µ-slide8 Ibidi, GmbH) functionalised with Poly-L-Lysine – 0.1% (P8920 Sigma Aldrich) and imaged in 0.4 ml/well Oxyrase-based STORM buffer (Phosphate Buffer Saline, 50mM Mercaptamine, 10 mM D-Lactate and 10 µl of a debris free SAE0010-5ml EC-Oxyrase Sigma).

These beads were imaged using a 100x 1.46 NA objective (*α* Plan-Apochromat 100×/1.46 NA Oil, Zeiss) and the Photometrics Prime 95B camera captured a FOV of 120 *µm* x 120 *µm*. Image Z-stacks were acquired over a range of -3 *µm* to +3 *µm* in 10 nm steps for 10 different FOVs of fluorescent beads such that in total approximately 1000 beads were imaged. The motorised XY stage was used to move laterally between different FOVs.

The training calibration data was acquired using a range of camera exposure times for the 10 different stacks, varying between either 300 ms, 500 ms or 1000 ms. The power at the sample was varied between settings of either 450 µW, 1.3 mW, and 2 mW. These low excitation powers were chosen as to minimise bleaching of the fluorescent beads during the acquisitions. The sample was excited at a wavelength of 635 nm.

### 2.3 Calibration data pre-processing

Starting from a full-FOV bead image stack, the most in-focus image frame was identified using the maximum pixel intensity as a function of axial displacement (Z). Individual beads were localised in this image using PICASSO^30^, and 3D image stacks for each bead were retrieved across the full FOV Z-stack. Similar sets of bead ROI Z-stacks from all full FOV image Z-stacks recorded were concatenated to produce a single training dataset.

Beads were excluded from the set of training data for any of the following issues: co-localisation with other beads within the cropped 15×15 pixel frame; poor signal-to-noise ratio (SNR, calculated as the ratio of maximum pixel intensity to mean pixel intensity for the bead’s image stack - a minimum value of 2 was used); or excessive noise in the Z or X/Y axes (measured by fitting a 1D or 2D skew Gaussian to the bead ROI Z-stacks in Z or X/Y respectively). The minimum SNR threshold and maximum noise thresholds were chosen through repeated training and evaluation of the model on test beads. Further tuning of the parameters was then informed by the reconstruction quality of nuclear pore samples.

The assignment of modelled Z-positions for each bead in the training data was key to removing bias from the final model, as two closely neighbouring beads with a similar PSF but different Z offsets could cause the model to learn an average of the two Z positions. The relative offset (*δZ*) of each bead in a FOV was calculated by finding the pixel-wise shift in Z which minimises a mean squared difference of pixel intensity values between any bead and a reference bead (chosen as the bead closest to the centre of the FOV). The overall *Z* = 0 defocus was selected using the frame in the reference bead stack with the largest image sharpness metric, and the relative *δZ* position of all other frames was calculated using the Z-step size of the acquired image stack (10 nm in this dataset).

The training dataset was formed of the 2D image, X/Y position, and Z position of each bead. Datapoints outside of the desired Z-range (±1 µm in our study) were removed from the dataset; this range can be modified to approximately match the microscope’s axial resolution (e.g. axial width of PSF). Beads were divided into training/validation/test sets and the training images were augmented to triple the number of training datapoints by adding Gaussian noise (*µ* = 0, *σ* = 0.005), randomly adjusting the brightness of the image (using a Keras RandomBrightness layer with a factor of 0.075^35^) and adding Poisson noise (*λ* = 226). These parameters were found using the hyper-parameter discussed in Section 2.4. Images of PSFs were resized to 64×64 pixels to allow the use of pre-trained weights (see Section 2.4). Pixel intensities were rescaled to a range of [0, 1] by dividing by the maximum possible pixel intensity. X/Y coordinates were rescaled to a range of [-1, 1].

### 2.4 Model architecture and training

The CNN model’s architecture (see Figure 2) was built around a MobileNet V3, which is a recent version of an efficient and fast CNN^32^. Pretrained ImageNet-21k^31^ weights were used to accelerate the training process. Future models could be trained from the weights shared in this paper to further reduce training time.

**FIGURE 2:**
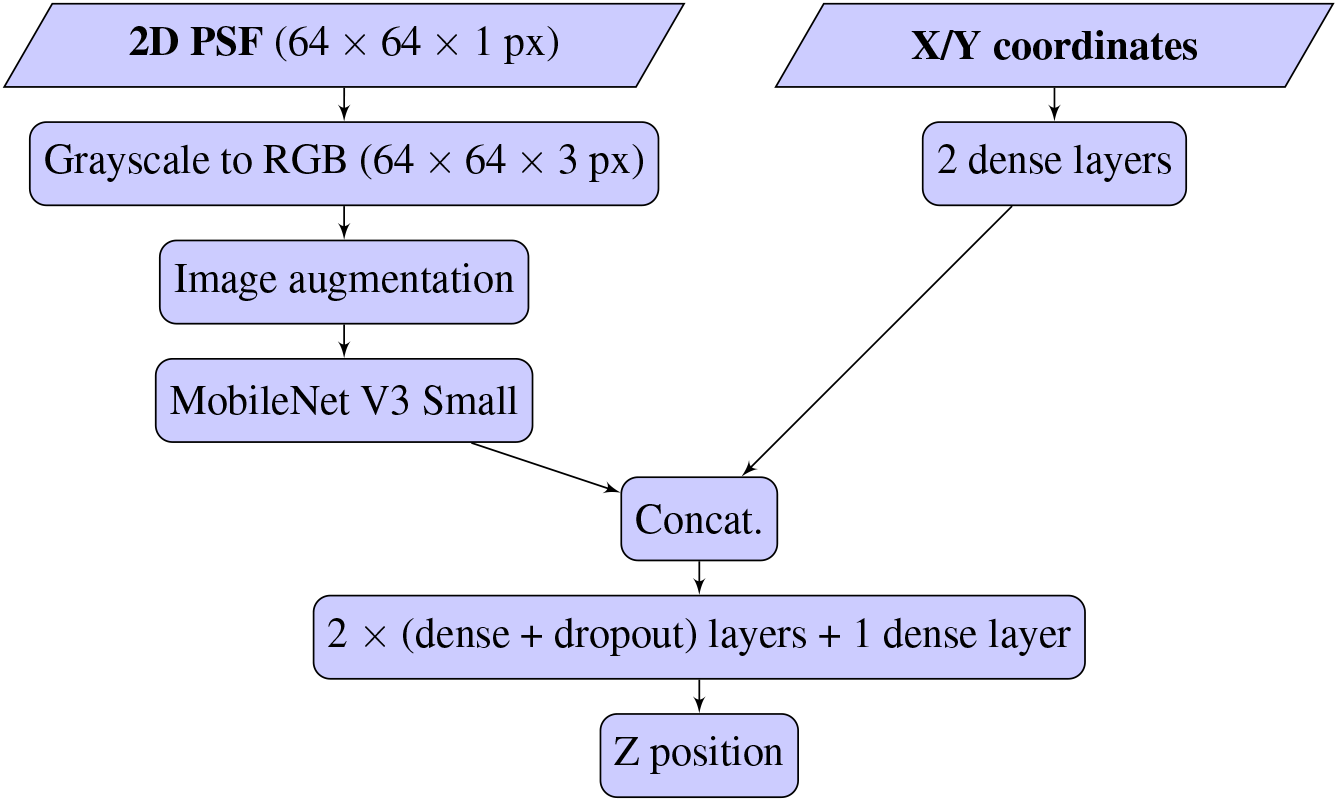
Model architecture including MobileNet V3 (small)^32^

Images of localisations were passed into the MobileNet V3 for feature extraction. X/Y coordinates were passed through two dense layers. Both input branches of the model were visible in Figure 2. The concatenated embedding of the image and coordinate branches of the network was then passed through a series of dropout and dense layers to output the predicted Z position.

The model was compiled with an AdamW optimizer^36^ and a mean squared error loss. Hyper-parameter optimisation from a Bayesian-guided search (as implemented in^37^) yielded specific augmentation parameters, dense layer sizes, learning rates and batch sizes. Trained models and associated training parameters are included in the Supporting Information. Callbacks were used to regularise the training; learning rate reduction was applied if the training loss stagnates, and an early stopping mechanism halts the training if the validation loss ceases to decrease.

The model was trained on an Ubuntu desktop equipped with an Intel Core i7-10700K CPU, a single NVIDIA RTX 3090 24GB graphic card and 128 GB of RAM, although this far exceeds the computational requirements of the small network size (3.4M parameters). Training our final model took approximately 1.5 hrs, which was significantly faster than for comparable methodologies trained from the same set of bead stacks (6 hrs for *DECODE* on the machine listed above, 14 hrs for *FD-DeepLoc* on a Windows machine with an AMD Ryzen Threadripper 3960X 24-Core 3.79 GHz Processor, 64.0 GB of RAM and a single NVIDIA RTX 3080 24GB graphic card). Note that whilst *DECODE* was trained on the same machine as *easyZloc, FD-DeepLoc* includes some Windows dependencies and was therefore trained on the aforementioned Windows machine of comparable hardware specifications. We consider that the difference between the computational hardware would not significantly impact the overall difference in training time.

### 2.5 Sample preparation and imaging

SNAPtag-labelled NUP96 nuclear pore proteins in the U-2OS-CRISPR-NUP96-SNAP cell line clone^38^ (CLS Cell lines Services GmbH) that we have labelled with BG-iFluor647^18^ were imaged using *easySTORM* implemented on the Axiovert 200 microscope described above. The preparation and imaging followed the protocols shared previously^10^. Briefly, cells were fixated for 1 minute with Paraformaldehyde 2.4% in TRB buffer (20 µM HEPES, 1 mM EGTA, 10 mM Potassium Acetate, 10 mM Sucrose), washed twice for 5 minutes each in Paraformaldehyde-free-TRB-Buffer, PFH quenched 5 minutes in freshly made 100 mM Ammonium Chloride in Phosphate Buffer saline (PBS), washed 3X in PBS, blocked for 10 minutes with Blocking buffer (3% Bovine Serum Albumin in PBS, 1µM Dithiothreitol), stained overnight with 1 µM SNAPtag in blocking buffer, and washed 3 times prior to imaging in Oxyrase buffer. The sample was incubated for 5 minutes with 1:1000 TetraSpeck beads in PBS followed by 3 washes with PBS to removed unbound beads to mark fiduciary markers. These SNAPtag-labelled nuclear pore samples were imaged on the Axiovert 200 Zeiss microscope and excited at 635 nm, using HiLo illumination^33^ for the SMLM acquisition. A power density of 1 kW/cm2 at the sample plane was used during the acquisition. 20000 frames were acquired with a camera exposure of 30 ms. Microtubules labelled with Docetaxel conjugated to Silicon Rhodamine (Sir-DNA, Spirochome) in NIH3T3 cells were also imaged using the same *easySTORM* microscope. Sample preparation followed the protocols of references^39,40^: cells were washed live at room temperature with a microtubule stabilizing buffer (PEMP buffer consisting of: 100mM PIPES pH6.8, 1mM EGTA, 2mM MgCl2, and 4% PEG 8000), permeabilised for 90 s with 0.5% Triton-X 100 in PEMP buffer, washed 4 times in PEMP Buffer, fixed in 0.2% glutaraldehyde in PEMP buffer for 15 minutes, and followed by treatment in 2 mg/ml Sodium Borohydride in PEMP buffer (NaBH4, dissolved immediately before use) for 15 minutes; samples were then washed 4 times in PEMP buffer, followed by incubation for 15 minutes with Docetaxel conjugated with Silicon Rhodamine (Sir-Tubulin Spirochome). To improve blinking for *dSTORM* imaging, samples were incubated at room temperature 5 minutes with freshly made NaBH4 (1 mg/ml) in PBS to modify fluorescent dye in a process known as reductive caging^41^. The sample was incubated for 5 minutes with 1:1000 TetraSpeck beads in PBS followed by 3 washes with PBS to removed unbound beads to mark fiduciary markers. This tubulin-labelled sample was imaged on the Axiovert 200 Zeiss microscope, using the same objective lens and camera as for the calibration bead stacks (see Section 2.3). The tubulin was also excited at 635 nm. The *easySTORM* dataset was acquired using a power density at the sample of 1.042 *kW*/*cm*^2^, using epi-illumination to illuminate the sample. 80000 frames were acquired at 30 ms exposure.

### 2.6 Reconstruction of 3D SMLM nuclear pore data

PICASSO^30^ was used for lateral single molecule localisation (X-Y) and lateral drift correction was applied using PICASSO via fiduciary markers (0.1 µM T7279 TetraSpeck Microspheres, Life Technologies). Pixel intensities were rescaled to a range of 0 to 1 by dividing by the maximum pixel intensity (65535, the maximum unsigned 16-bit integer), and X/Y coordinates were rescaled to a range of -1 to 1 in accordance with the FOV of the training data. The pre-processed data was then localised in Z using our trained CNN model to produce a set of 3D localisations. Drift correction in Z was implemented by calculating the mean Z localisation of 500 sequential image frames and fitting a smoothed cubic spline to the mean Z position for all localisations in the frame. The Z positions for localisations in each frame were then corrected by subtracting any offset relative to the mean Z localisation over time.

Nuclear pores were picked in X/Y using PICASSO’s Render GUI and analysed individually. Each group of localisations was modelled using a kernel density estimation (KDE) with a bandwidth of 15 nm, selected to image structures on a scale of 50 nm (as expected for the Nup96 protein rings) without over-fitting to noise. KDEs smooth the data to minimise the impact of noise and highlight the underlying spatial structure of data. A KDE was applied by evaluating a Gaussian kernel over the range of Z values of localisations in each nuclear pore. In order to quantify the success at resolving nuclear pore structure, the reconstructed data was automatically scanned to identify reconstructed features (i.e., peaks observed with an applied Gaussian KDE, bandwidth = 15 nm, min peak prominence of 0.0001 localisations / *nm*^2^) that could correspond to a nuclear pore’s two layers of labelled Nup96 proteins. A nuclear pore was considered to be “successfully” imaged if the two more prominent peaks are axially separated by 40 to 60 nm, following the criteria in^21^. The visual contrast of the reconstructed image of the nuclear pore layers was enhanced by applying a minimum threshold for the spatial density. This threshold was calculated as the mean of the mean peak height and the local minima.

## 3 RESULTS

*easyZloc* was trained on a bead dataset obtained using the methodology in Section 2.2. The model was evaluated on bead data withheld from the calibration data. Figure 3 illustrates how the *easyZloc* model correctly learned to map field dependent aberrations to the Z position of the bead. The resulting Z localisation accuracy is comparable to those reported by alternative methodologies on the same calibration dataset (Root Mean Squared Error of Z localisation, easyZloc: 120 nm, *DECODE*: 148 nm, *FD-DeepLoc*: 156 nm).

**FIGURE 3:**
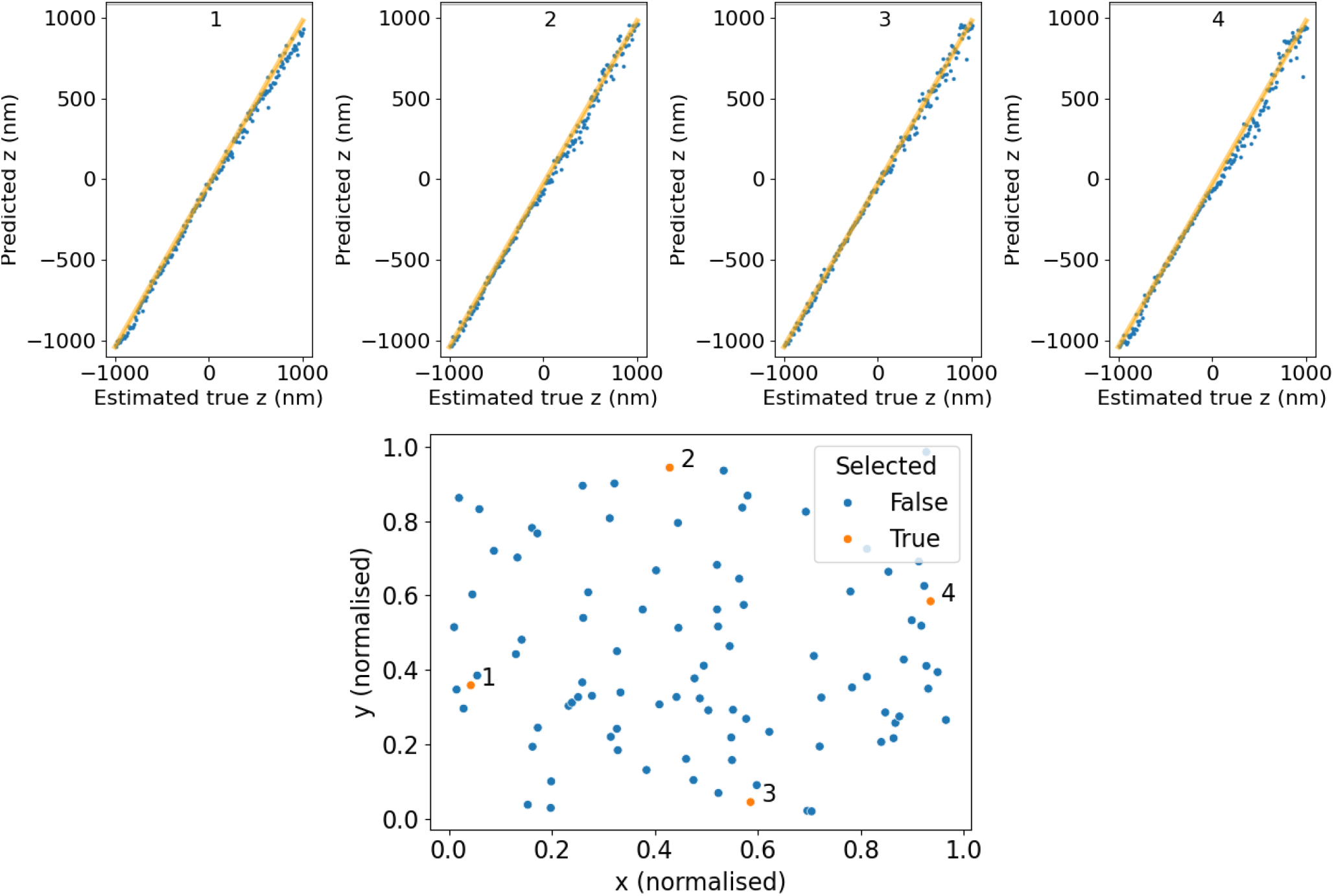
Performance of *easyZloc* on test beads. Top: each plot (1-4) was produced from a bead within the Z stack of the training data. The position in Z of each slice in the Z stack was estimated from the peak illumination of the stack and the Z-step of the microscope system (10 nm). The model estimates positions for each slice of the Z stack, and we see a linear correlation (orange line) with the stage movement of the microscope. The model achieves a root mean squared error of 40.93 nm, 39.46 nm, 36.36 nm and 55.35 nm respectively on these relatively noise-free beads. Bottom: orange dots (1-4) show the position of test beads in the FOV, demonstrating the model has learned to map laterally varying aberrations.

The values for *DECODE* and *FD-DeepLoc* were automatically generated by the respective methodology’s model training software. We note, however, the different training and test data for each methodology. *easyZloc* directly utilised empirical image Z-stack image data of 20 nm Polystyrene bead (F8782-Invitrogen FluoSpheres) fluorescent beads, whereas *DECODE* and *FD-DeepLoc* were applied using a physical model of the PSF constructed from these 20 nm bead image data that was then used to simulate images of the PSF which were then used for training.

We note that while *DECODE* and *FD-DeepLoc* can function with any modellable PSF modality, they were designed for and demonstrated on microscopes with additional induced astigmatism. The optical system used in our implementation is a more challenging case for Z localisation due to less pronounced changes in the PSF over Z; *DECODE* and *FD-DeepLoc* may therefore provide a lower axial resolution than they would if applied with an engineered PSF.

The trained models were then applied to *easySTORM* SMLM data of Nup96 nuclear pore proteins in the U-2OS-CRISPR-NUP96-SNAP cell line clone^38^ (CLS Cell lines Services GmbH) that we have labelled with BG-iFluor647^18^. While *DECODE* and *FD-DeepLoc* directly yield 3D localisation tables from the SMLM data, with our approach the SMLM data was first analysed using PICASSO^30^ to obtain the 2D localisation table, and then Z localisation was undertaken using *easyZloc*.

Nuclear pore complexes were extracted from reconstructions of the dataset by each method to explore the relative success rates with which Nup96 layers of the nuclear pore could be distinguished (we used this as an arbitrary definition of a ‘successful’ reconstruction, as discussed in Section 2.6). Exemplar reconstructed nuclear pore images are shown in Figure 4, and additional examples can be seen in the Supplementary Information in Figure 8.

**FIGURE 4:**
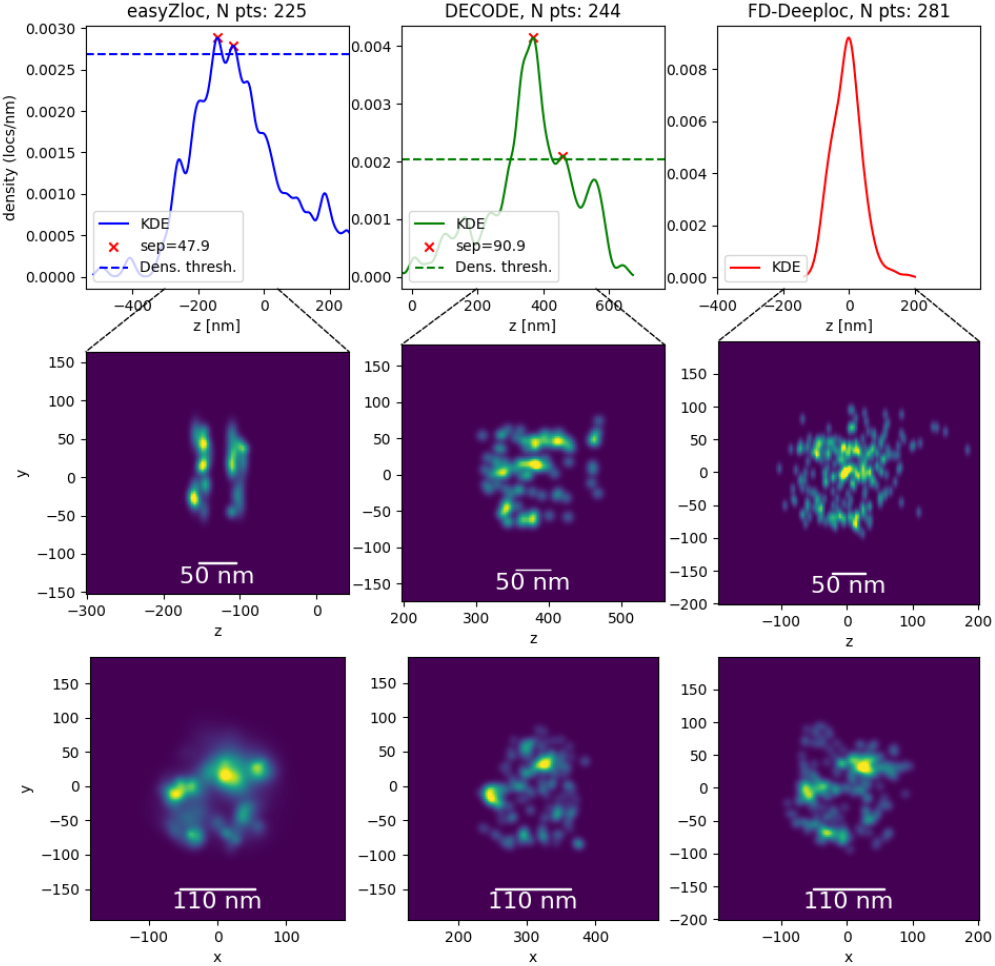
Reconstructions of Nup96 proteins in a nuclear pore imaged with a 100x 1.46NA objective lens and analysed using *easyZloc, DECODE* and *FD-DeepLoc*. Rendered X/Z images are thresholded by a minimum spatial density to increase the visual contrast of the image, as indicated by the dotted line of each histogram. The threshold density is set automatically following the procedure detailed in Section 2.6. The X/Y view was not thresholded has the contrast was already sufficient to identify the NPC structure. Additional comparative examples of nuclear pore reconstructions can be found in the Supplementary Information in Figure 8, where three “successful” reconstructions of nuclear pores for each methodology are displayed with the corresponding reconstruction of the same data using the other methodologies. Also provided in the Supplementary Information is a video similarly presenting all the nuclear pores that were “successfully” reconstructed by at least one methodology.

Figure 5 plots the number of ‘successful’ 3D nuclear pore images reconstructed from the same *easySTORM* SMLM data set using *easyZloc, DECODE* and *FD-DeepLoc*, plotted as a function of the KDE smoothing parameter (see Section 2.6). We note that (MobileNet v3-based) *easyZloc* performs at least as well for KDE bandwidth values above 7 and functions up to a KDE bandwidth >20 – at a significantly lower computational cost (see Section 2.4).

**FIGURE 5:**
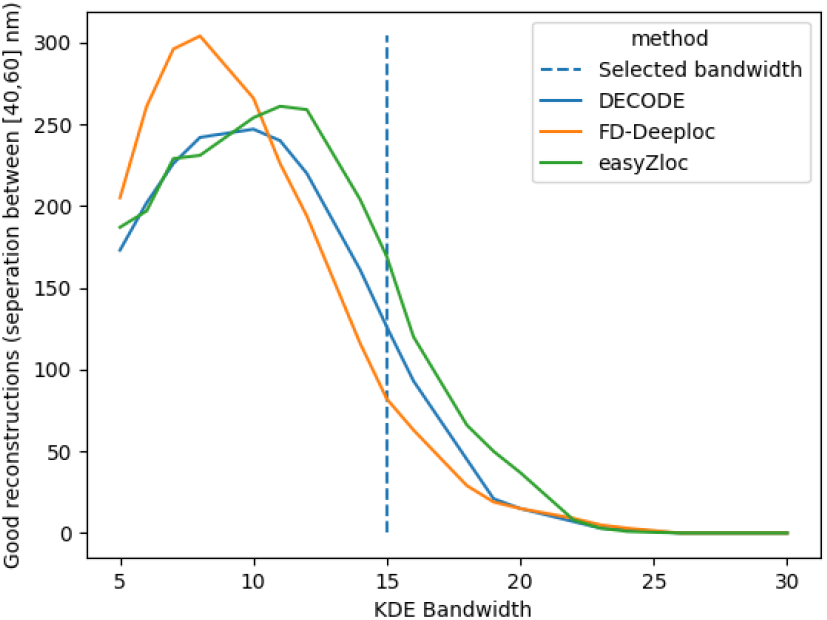
Comparison of the number of “successfully” resolved nuclear pores for each Z localisation methodology applied to the same dataset. The criteria to assess nuclear pore reconstructions is detailed in Section 2.6.

The choice of a KDE bandwidth affects the number of nuclear pores reconstructed. An excessively large bandwidth would blur the data, causing the nuclear pore’s layers to be indistinguishable, whilst a small bandwidth can lead to overfitting of noise. Standard bandwidth tuning methods, as discussed in^42^, resulted in unresolvable reconstructions due to over-smoothing of the data. We note that KDEs have been utilised in previous work^21^ although the methodology for selecting the value of the KDE bandwidth was not discussed.

The depth colour-coded 3D SMLM reconstruction in Figure 6 illustrates the depth range of *easyZloc* applied to the same *easySTORM* SMLM data (following 2D localisation using PICASSO) of Nup96 nuclear pore proteins within the cellular nuclear envelopes, in a large (>100 µm) FOV. The data processing for this entire FOV required less than 10 minutes of compute on a Linux computer (hardware details are listed in Section 2.4). The equivalent reconstructions using *DECODE* required significantly more computational time (2.74 hrs) on the same machine as *easyZloc*, while *FD-DeepLoc* required 0.3 hrs on a similar machine using Windows (see hardware details in Section 2.4).

**FIGURE 6:**
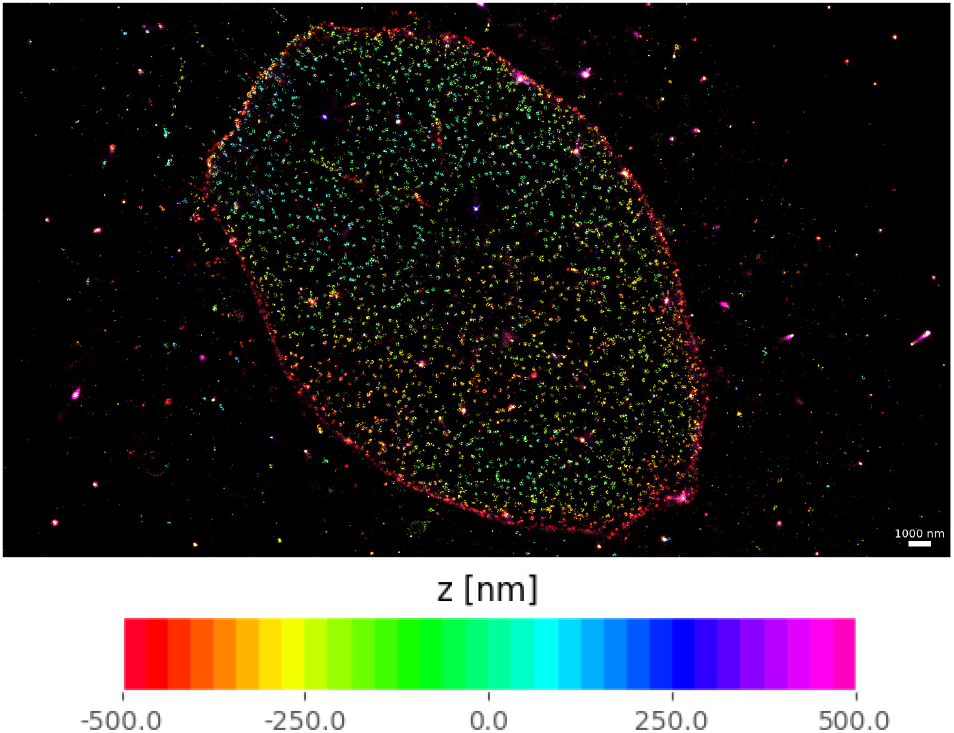
3D reconstruction of a cell nucleus imaged with a 1.46 NA objective lens (*α* Plan-Apochromat 100×/1.46 NA Oil, Zeiss), coloured by depth, as produced by *easyZloc*. Scalebar length is 1µm.

Figure 7 shows a 3D SMLM reconstruction of *easySTORM* image data (following 2D localisation using PICASSO) of tubulin in NIH3T3 cells labelled with Sir Tubulin acquired on the same epifluorescence microscope. Here *easyZloc* provides depth information over 2 µm.

**FIGURE 7:**
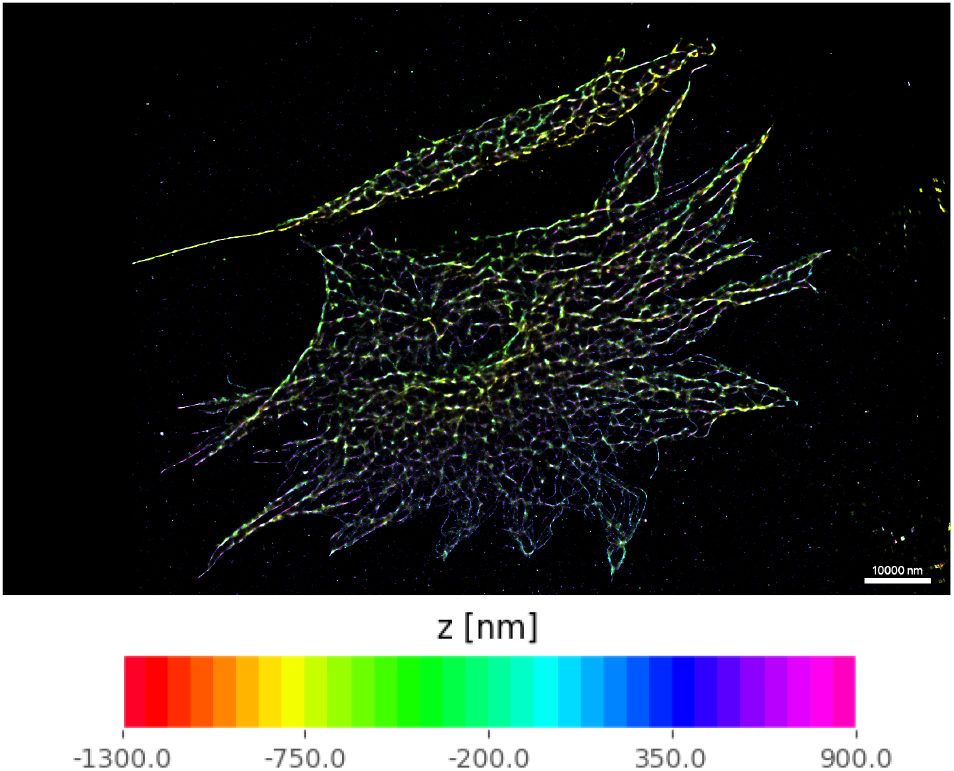
3D SMLM reconstruction of tubulin in NIH3T3 cells imaged with a 1.46 NA objective lens (*α* Plan-Apochromat 100×/1.46 NA Oil, Zeiss) objective lens using *easyZloc* following 2D localization using PICASSO. Scale bar is 10 µm.

## 4 DISCUSSION

We have presented a fast and lightweight machine learning-based methodology for axial localisation of SMLM data, *easyZloc*, which can be used in standard epifluorescence microscopes without specifically engineering an axially varying PSF. Requiring only the additional acquisition of a calibration Z-stack of images of sub-resolution fluorescent beads, *easyZloc* can be implemented with existing 2D SMLM workflows using established 2D localization software. Since *easyZloc* performs only a subtask within the overall SMLM localisation task - sequentially identifying the defocus of individual localisations from an extracted FOV centred on that localised emitter - it is less computationally demanding than 3D localisation software tools that identify multiple localisations in their inputs and determine the full 3D localisation of the SMLM dataset. Furthermore, the use of empirical training data circumvents the need for mathematical modelling of the PSF, allowing for the deep learning model to readily account for lateral variation in the microscope. Finally, the use of the latest lightweight deep learning architecture (MobileNet V3 small) means that the methodology requires comparatively modest computational resources and a short execution time. We thus believe this approach has broad potential to widen access to 3D SMLM and may be useful when scaling to higher throughput in automated SMLM instruments.

We note that the choice of the core CNN architecture, MobileNet V3 (small version), was made after considerable hyper-parameter exploration using the nuclear pore Nup96 cell line as a test sample. Vision Transformers^43^ were originally used but suffered from over-fitting issues, whilst not improving the frequency or clarity of nuclear pore reconstructions. ResNets^44^ and VGG^45^ architectures presented similar issues. We surmise that these more powerful methodologies may surpass *easyZloc* in speed and precision if given a larger quantity or variety of training data, and/or when implemented on more powerful deep learning architectures.

## Supporting information

Video of nuclear pore reconstructions

## ACKNOWLEDGMENTS

We acknowledge funding from the Cancer Research UK (A29368) and from the Chan Zuckerberg Initiative DAF, an advised fund of the Silicon Valley Community Foundation (grant 2021-234618, 2023-321240). Miguel Boland acknowledges a PhD studentship supported by the Wellcome Trust grant 203799/Z/16/Z.

## DATA AVAILABILITY STATEMENT

Data and trained models are available upon request. The codebase supporting this work is available at https://github.com/mb1069/easyZloc.

## SUPPORTING INFORMATION

Figure 8 presents three “successful” nuclear pore reconstructions for each Z localisation methodology (*easyZloc, DECODE, FD-DeepLoc* along with corresponding reconstructions of the same data using the other methodologies

**FIGURE 8:**
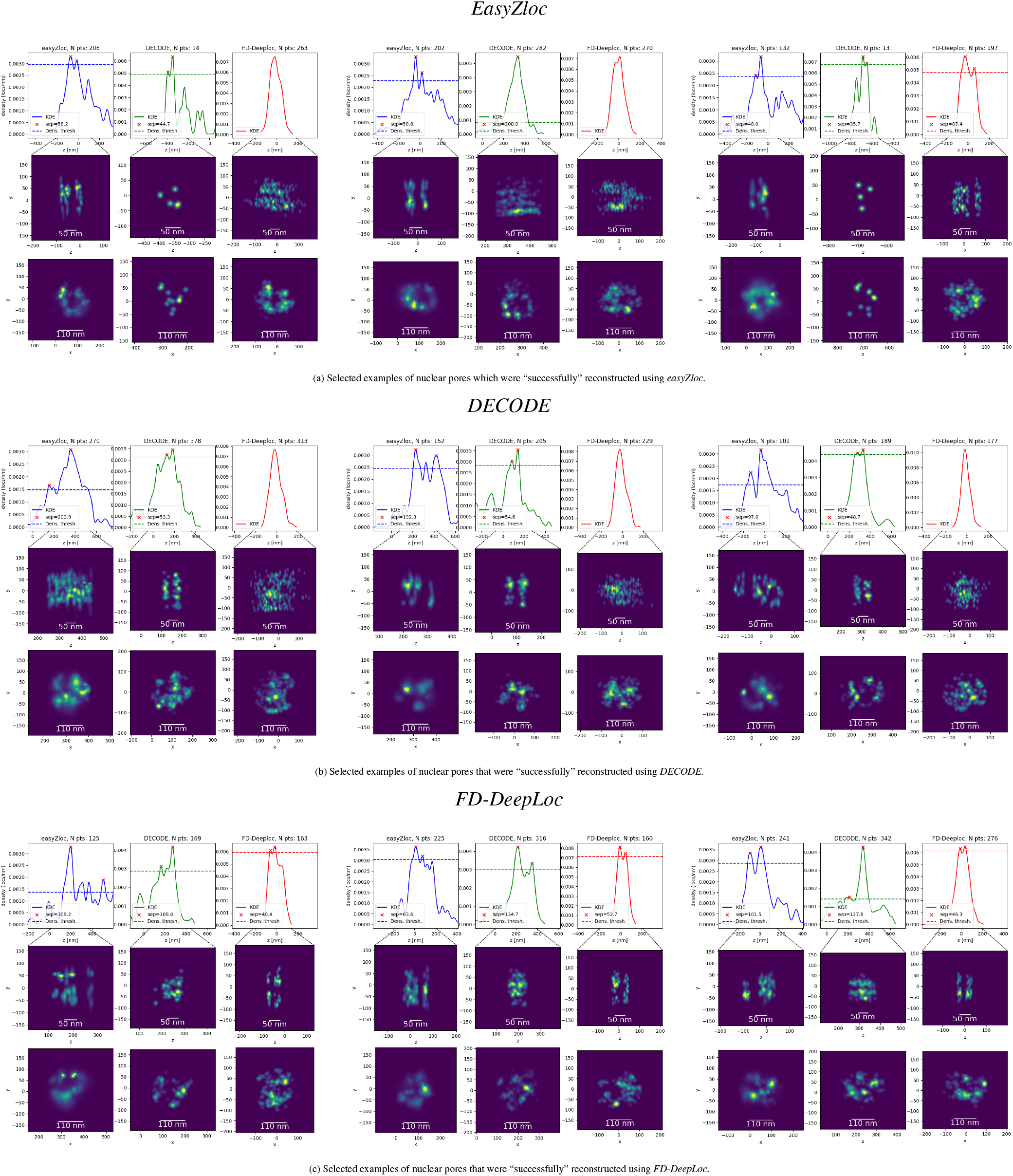
Additional examples of nuclear pore reconstructions

Supplementary Video 1 presents a comparison of reconstructions of all the nuclear pores which were “successfully” reconstructed by at least one Z localisation methodology.

